# Anomalous invasion dynamics due to dispersal polymorphism and dispersal-reproduction trade-offs

**DOI:** 10.1101/2020.02.19.956425

**Authors:** Vincent A. Keenan, Stephen J. Cornell

**Author notes:** Department of Mathematics and Statistics, University of Strathclyde, Livingstone tower, 26 Richmond Street, Glasgow G1 1XH, Scotland (Vincent A. Keenan). Correspondence should be sent to: Dr. Stephen Cornell, Institute of Integrative Biology, University of Liverpool, Crown Street, Liverpool L69 7ZB, UK. **Author contributions:** Both authors designed and performed the study, and both authors wrote the manuscript. **Data Accessibility:** Should the manuscript be accepted, the data supporting the conclusions will be archived in Figshare.

## Abstract

Dispersal polymorphism and mutation play significant roles during biological invasions, potentially leading to evolution and complex behaviour such as accelerating or decelerating invasion fronts. However, life history theory predicts that reproductive fitness — another key determinant of invasion dynamics – may be lower for more dispersive strains. Here, we use a mathematical model to show that unexpected invasion dynamics emerge from the combination of heritable dispersal polymorphism, dispersal-fitness trade-offs, and mutation between strains. We show that the invasion dynamics are determined by the trade-off relationship between dispersal and population growth rates of the constituent strains. We find that invasion dynamics can be “anomalous” (i.e. faster than any of the strains in isolation), but that the ultimate invasion speed is determined by the traits of at most two strains. The model is simple but generic, so we expect the predictions to apply to a wide range of ecological, evolutionary or epidemiological invasions.

## Introduction

Evolution can significantly affect the dynamics of biological invasions and range expansion (Hallatschek and Nelson, 2009; Kubisch et al., 2013; Hodgins et al., 2018). Deleterious mutations can accumulate during invasions, leading to a deceleration in the rate of advance (Peischl et al., 2015), but evolutionary processes can also facilitate invasions. Dispersal evolution has attracted particular interest, as higher dispersal ability confers a clear advantage when colonising new habitat (Kubisch et al., 2013). Empirical data suggest that newly established populations contain individuals with elevated dispersal capabilities (Simmons and Thomas, 2004; Fronhofer and Altermatt, 2015), and accelerating cane toad invasions are accompanied by an evolution of dispersal-related traits such as leg length (Phillips et al., 2006) and dispersive behaviour (Lindstrom et al., 2013). The tendency of more dispersive strains to be found at the vanguard of an invasion has been dubbed “spatial sorting” (Shine et al., 2011), a form of dispersal evolution that plays out in space as well as time (Phillips and Perkins, 2019).

However, invasion dynamics depend on population growth as well as dispersal (Fisher, 1937; Skellam, 1951). A higher investment in dispersal ability can imply a lower investment in traits related to reproduction (Saastamoinen et al., 2017), so an increase in dispersal ability does not necessarily imply a faster invasion. Moreover, a sufficiently strong trade-off could disrupt strict spatial sorting — a hypothetical example would be a highly dispersive but infertile strain, which would not advance the invasion. Simulation studies show that the rate of spread can accelerate in the presence of a dispersal-fecundity trade-off (Burton et al., 2010), but to predict the general conditions under which different invasion scenarios (e.g. spatial sorting, constant or accelerating speed, etc.) occur requires a mathematical theory. Several studies have shown that invasion speed for a polymorphic species with a trade-off is determined by the strain which would invade most quickly on its own (Osnas et al., 2015; Phillips and Perkins, 2019; Deforet et al., 2019), but those studies omit mutation — which can plays a key role in invasion dynamics by creating interactions between strains. Elliott and Cornell (2012) showed that, if a species comprises two strains (one a superior disperser, and one more fecund), with infrequent mutation between strains, it can invade at a significantly faster, “anomalous”, speed than would be predicted for either strain in isolation. However it is not clear how to generalise this result to a more realistic species consisting of multiple strains. For example, it is not obvious whether the invasion dynamics is determined by all of the strains or just a few, or how the invasion dynamics could be predicted for a species with a very large number of strains. Since both dispersal and reproductive fitness are beneficial, invasions could potentially promote a mixed strategy or even evolutionary branching (Weigang and Kisdi, 2015; Laroche et al., 2016). Here we define “reproductive fitness” as per capita growth rates at low density and use this definition throughout the article.

Here, we develop a general theory for invasions by species with dispersal polymorphism, a dispersal-fitness trade-off, and mutation between strains. We show that the shape of the trade-off curve critically determines whether the invasion speed continues to accelerate or approaches an asymptote. The trade-off curve also determines whether the invasion speed equals that of one of the constituent strains, or whether the speed is anomalous, i.e. is faster than any single strain on its own. Surprisingly, in all cases, we find that the asymptotic speed is determined by the traits of at most two of the constituent strains. We find that strong effects of spatial sorting are not realised in all cases — the most dispersive strains do not necessarily lead the invasion — and that the invasion speed is not necessarily determined by the most dispersive or the most fecund strain, or the one which, in isolation, would invade at the fastest speed. Our model is simple but generic but requires large population sizes, so is most likely to apply to microbial or microbial systems such as infectious diseases.

## Methods

We consider an asexual haploid species consisting of *N* strains (i.e. genotypes), where each strain has a distinct growth rate and dispersal phenotype, with the possibility of mutation between strains at birth (derivation can be found in Supplementary Information Appendix S1). We first consider a deterministic model in continuous space and time, for which we can derive exact results. We assume density-dependent competition between strains and that dispersal can be approximated by diffusion, and model the dynamics using the following spatial Lotka-Volterra model

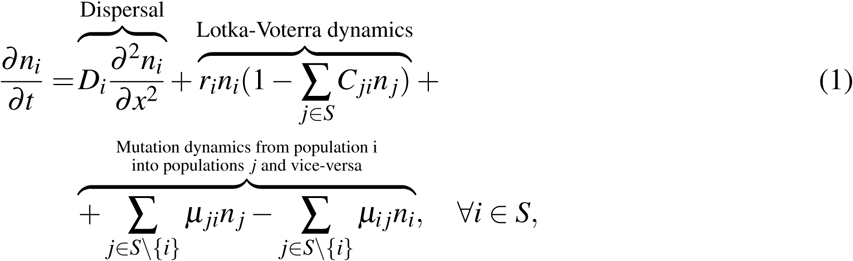

where *S* = {1, 2, …, *N*} is the set of all strains, *D*_*i*_ and *r*_*i*_ are respectively the diffusion constant and population growth rate of the *i*th strain, *n*_*i*_ is the density of the *i*th strain, *C*_*ji*_ is the competitive effect of strain *j* on *i*, and *µ*_*ij*_ the mutation rate from strain *i* to strain *j*. This extends the investigation of Elliott and Cornell from two strains to a general degree of population polymorphism (Elliott and Cornell, 2012). Equations including diffusion, Lotka-Volterra competition, and mutation between strains have been considered before (Dockery et al., 1998; Bouin and Calvez, 2014; Girardin, 2018), but explicit results for the case where both *D*_*i*_ and *r*_*i*_ differ among strains have not previously been computed for *N >* 2. Provided it is possible for any strain to mutate, after enough generations, into any other strain (i.e., provided the species does not comprise sub-species between which mutation is not possible) there is a single stable (spatially uniform) equilibrium.

We consider an invasion, where half of space is initially empty and the other half is occupied by the species at equilibrium. For this class of partial differential equations, Girardin (2018) has proven that the solutions to equations of the form eqn. (1) under these initial conditions are travelling waves of the form *n*_*i*_ = *N*_*i*_(*x − c*^*\**^*t*), where *c*^*\**^ is obtained by the solution of an eigenvalue problem. Using Theorem 1.7 of Girardin (2018), the spreading speed for eqn. (1) is given by

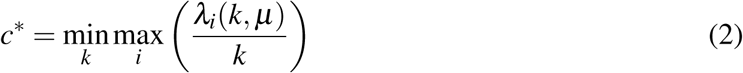

where *λ*_*i*_(*k, µ*), *i* ∈ *S* are the eigenvalues of the matrix

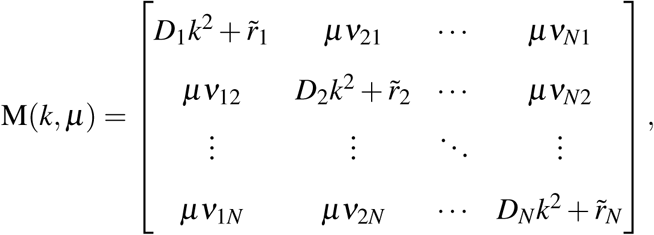

where 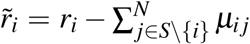 and we have defined 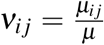 so that we can take the limit *µ →* 0 while keeping *ν*_*ij*_ = *O*(1). This same expression can be obtained by linearising eqn. (1) around the unstable equilibrium, using the ansatz *n*_*i*_ *∼* exp(*λt − kx*), and minimising the speed *c* = *λ/k* over *k*. However, until Girardin’s proof it was not known that the speed for the linearised equation would give the exact spreading speed for nonlinear PDEs of the form (1).

While this eigenvalue cannot be calculated in closed form for general *N*, we compute the limiting value of this wave speed when the mutation rate approaches zero, and confirm these results with numerical simulations. We expect this to be a good approximation for real species, where offspring usually inherit their parent’s traits and mutations between strains only happen occasionally (i.e. as *µ →* 0). When *µ* = 0 all of the off diagonal elements are zero so the eigenvalues equal

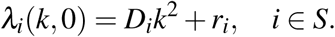

Taking the limit *µ →* 0 in eq. (2)

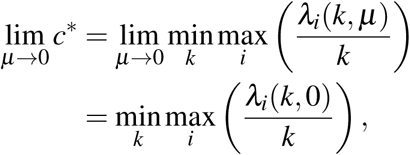

In the second line of the preceding equation, we have swapped the order of the limit *µ →* 0 with the minimum and maximum; this is justified because the eigenvalues of a matrix depend continuously on its elements, so the eigenvalues of *M* converge uniformly to *λ*_*i*_(*k, µ*) as *µ →* 0. We can therefore find the limiting value of the invasion speed *c*^*\**^ as *µ →* 0 by plotting the curves 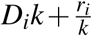 as a function of *k*, selecting the upper envelope of these curves, and then finding the value of *k* for the lowest point on this upper envelope. More details of each of the steps in the calculation, and a graphical illustration, can be found in Supplementary information Appendix S1.

It is known that demographic stochasticity can affect anomalous speeds in dimorphic species (Elliott and Cornell, 2013), so to test the robustness of our results we also ran simulations of a stochastic Beverton-Holt model. This is based on the following deterministic model:

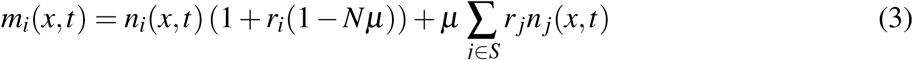

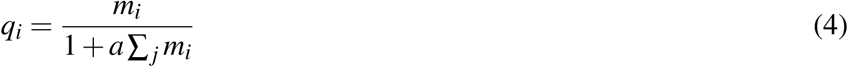

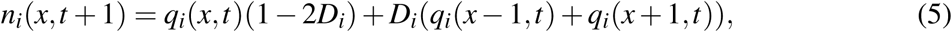

where eqn. (3) represents reproduction with mutation, eqn. (4) density-dependent mortality, and eqn. (5) dispersal, except Poisson and Multinomial pseudorandom number generators replace eqns. (4) and (5) (more details in the Supplementary information Appendix S1). Here, *r*_*i*_ and *D*_*i*_ again represent population growth rate and dispersal ability, and *µ* represents mutation. The parameter *a* sets the scale of density dependence, so that the stable equilibrium density is proportional to 1*/a*. We expect that the stochastic and deterministic version of the model will be similar when *a* is very small.

## Results

We find that anomalous invasion speeds are possible in the *N*-strain case, just as was found in the 2-strain case (Fig 1). When the mutation rate is very small but nonzero (i.e. in the limit *µ →* 0), the speed of invasion by any *N*-strain species is obtained by calculating the maximum of 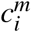 and 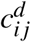 in the following expressions:

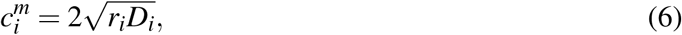

for all *i*, and

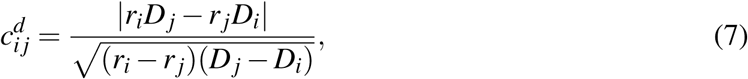

for all pairs *i* and *j* such that

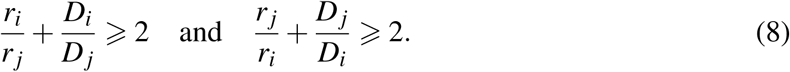

**Figure 1:**
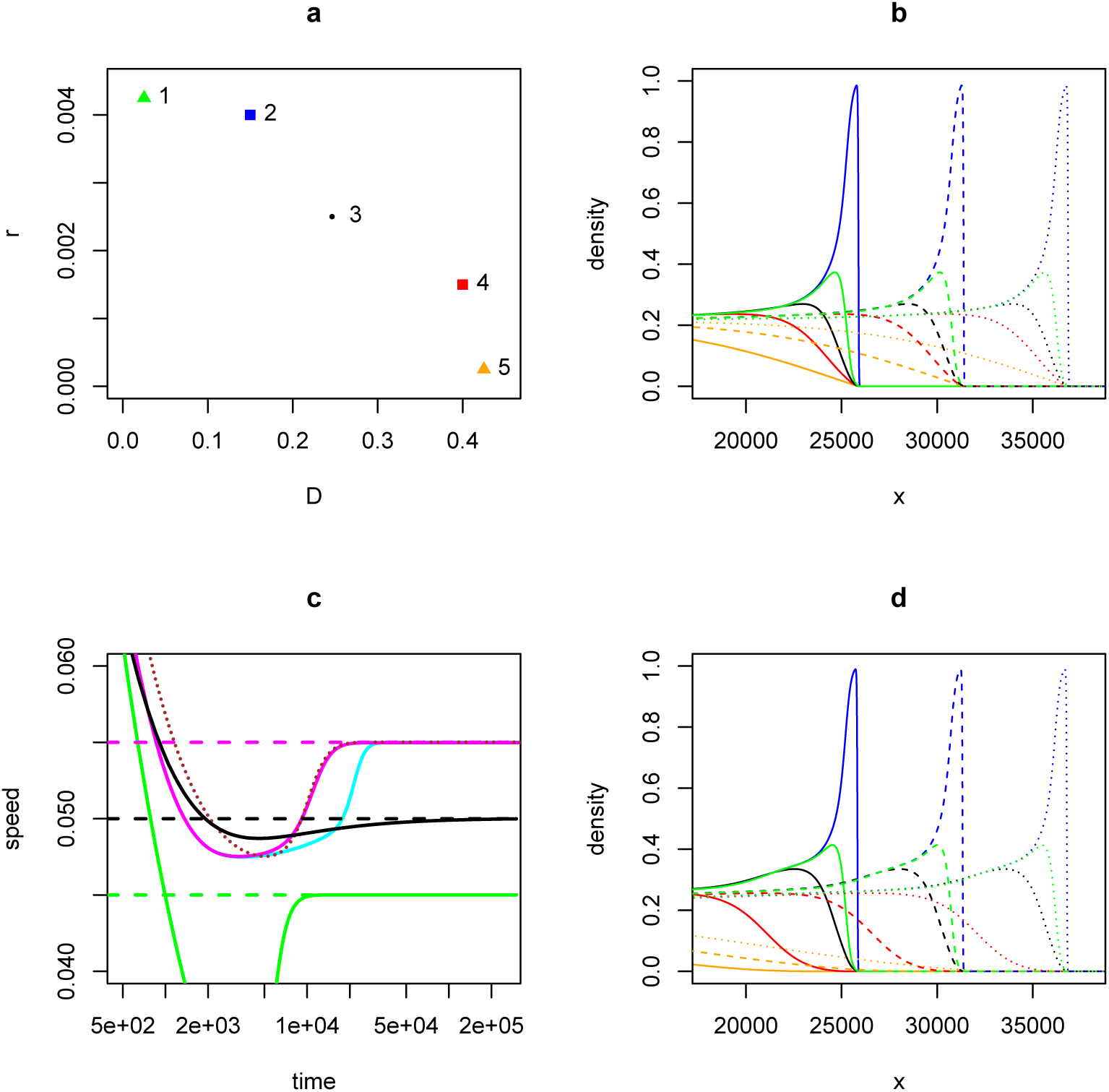
Anomalous invasions by polymorphic species into unoccupied habitat, where all strains can coexist at equilibrium, from numerical solutions of eqn. 1. Panel a: trait values for 5 strains used in this figure. (*D*_*i*_, *r*_*i*_) = (0.025, 0.00425), (0.15, 0.004), (0.25, 0.0025), (0.4, 0.0015), (0.425, 0.00025) for strains *i* = 1, 2, 3, 4 and 5 respectively. Squares denote the “vanguard” strains (2 and 4) and triangles the “extreme” strains with the highest growth rate (strain 1) and highest dispersal ability (strain 5) respectively. Strain 3 (circle) has the highest monomorphic speed. Colours correspond to those used in panels b and d (green, blue, black, red, and orange for *i* = 1, 2, 3, 4, 5 respectively). Panels b and d: Population density as a function of spatial coordinate *x* during invasion by a species consisting of all 5 strains, displaying travelling waves, at times *t* = 4 *×* 10^5^(solid curves), 5 *×* 10^5^(dashed curves), 6 *×* 10^5^(dotted curves). Mutation is universal among strains (*ν*_*ij*_ = 1∀*i, j*) in panel b, but in Panel d is only nonzero between neighbouring strains (*ν*_*ij*_ = 1 if |*i - j*| = 1, 0 otherwise). Panel c: Speed of position of front for species consisting of different combinations of strains. Magenta: all 5 strains present, universal mutation. Cyan: all 5 strains present, neighbouring-strain mutations only. Brown dotted curve: vanguard strains (2 and 4) only. Black: strain 3 (fastest monomorphic speed) only. Green: extreme strains (1 and 5) only. Dashed horizontal lines are theoretical predictions for 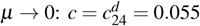 (magenta); 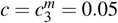 (black); 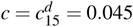 (green). In all cases mutation rate *µ* = 10^−6^, and competition coefficients are *C*_*i j*_ = 1 for *i* = *j*, 0.9 otherwise.

Here, 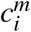 is the “monomorphic speed” at which a species consisting solely of strain *i* would invade. 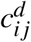 is the “dimorphic speed” at which a species consisting solely of strains *i* and *j* would invade, provided conditions (8) are met (Elliott and Cornell, 2012). Conditions (8) imply that valid dimorphic speeds only exist for pairs of strains whose dispersal and growth rates differ sufficiently – a graphical representation can be seen in Elliott and Cornell (2012) Fig 1. If the largest valid 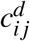 is greater than the largest 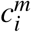, then the invasion is more rapid than that for any of the constituent strains in isolation, and the invasion is said to be “anomalous” (Elliott and Cornell, 2012).

Our analysis shows that an *N*-strain species therefore invades at the same speed as if it consisted of only one or two of its constituent strains — the traits of the other strains do not affect the invasion speed. This is illustrated by simulation in Fig 1 for a species consisting of 5 strains. Parameters are chosen (Fig. 1a) so that the strains that are predicted to determine the invasion speed (2 and 4) are not the strain with the highest monomorphic speed (strain 3) nor the strains with the highest population growth rates or dispersal (strains 1 and 5). Fig 1c shows that, after a transient, the invasion for a 5-strain species (magenta curve) advances at the same speed as a species containing only strains 2 and 4 (brown dotted curve), which are the strains with the highest dimorphic speed 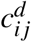 from eqn. (7). This is faster than either strains 2 or 4 in isolation (not shown), or the strain with the fastest monomorphic speed in isolation (black curve). Thus, while anomalous invasion dynamics implies a synergy between two strains (benefiting from the growth rate of a more fecund strain and the dispersal ability of a more dispersive one), this synergy does not extend beyond more than two strains. Moreover, one might expect that the “vanguard” strains (i.e., the two strains whose traits determine the anomalous speed) would be the most dispersive and the most fecund, so that the population as a whole benefits from the highest diffusion constant and the highest population growth rate. However, it turns out that this need not be the case: a species consisting of strains 1 and 5, which have respectively the highest fecundity and dispersal ability, invades more slowly (green curve in Fig. 1c). Note from Fig. 1b that the invasion shows weak effects of spatial sorting: the invasion is led by the second least dispersive strain (blue curve), and the strains with the highest monomorphic speed (black) and dispersal (orange curve) trail behind. In all cases, the long-term invasion speeds closely match the predictions for small mutation rate from eqns. (6) and (7) (horizontal dashed lines in Panel 1d).

The analysis predicts that the invasion speed is determined by the dynamics at low densities, and is therefore independent of the competition coefficients *C*_*ji*_. We therefore obtain the same anomalous speeds, whether or not the strains coexist within the range core (i.e. at the stable equilibrium). In Fig 2a, the invasion speed of a species consisting of stains 2, 3, and 4 (magenta) is given by the anomalous speed for strains 2 and 4 (brown dotted curve), even though these are both out-competed by strain 3 in the core range. This shows that strains that are very rare in the range core of the species can still determine the invasion dynamics of the species. Strain 2, which has the highest population growth rate, leads the front with high density (Fig 2b, blue curve). Strain 4, which has the highest dispersal ability, has a higher density in the wake of the front than at equilibrium but lower density than the other two strains (Fig 2b, red curve). Nevertheless, removing either strain 2 or 4 slows the invasion (Fig 2a, red and blue curves), to a slower but still anomalous speed (cyan horizontal dashed line), which is quicker than the invasion speed for strain 3 alone (black), which in turn is quicker than either vanguard strains 2 or 4 in isolation (not shown).

**Figure 2:**
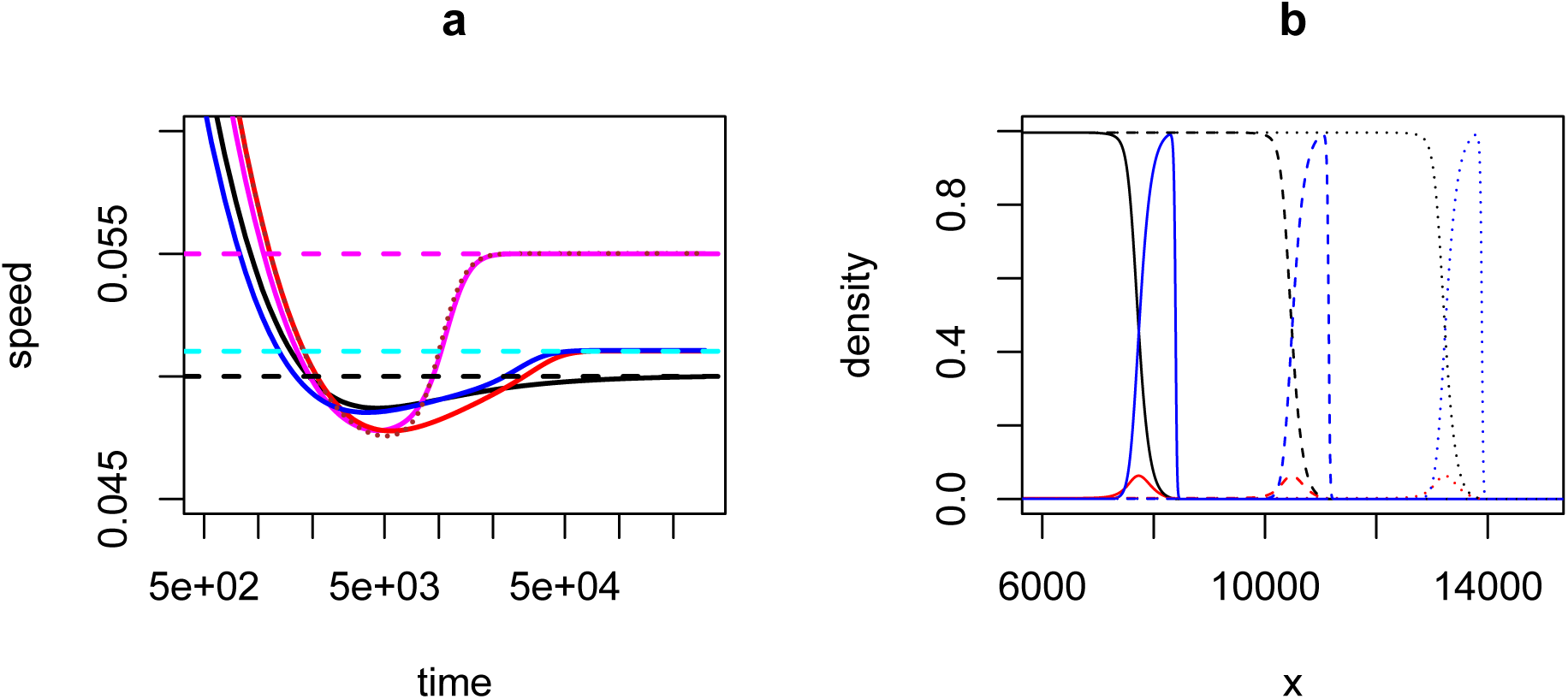
Strains that are outcompeted at equilibrium can still generate anomalous invasion speeds. Strains are numbered as in Fig. 1q, but competition coefficients are such that strain 3 outcompetes strains 2 and 4 at equilibrium (*C*_*ji*_ = 0.8 for *i* = 3 & *j* = 2 or 4; *C*_*ji*_ = 1.25 for *j* = 3 & *i* = 2 or 4). Panel a: Invasion speeds as a function of time. Colours denote different combinations of strains being present in the species: strains 2,3,4 with universal mutation (magenta); strains 2 and 4 (brown dotted); strains 2 and 3 (blue); strains 3 and 4 (red); strain 3 only (black). Dashed horizontal lines are theoretical predictions for 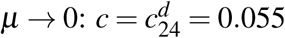 (magenta); 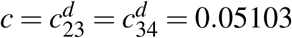 (cyan); 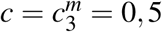 (black). Panel b: Population density as a function of spatial coordinate *x*, at times *t* = 10^5^(solid curves), 1.5 *×* 10^5^(dashed curves). 2 *×* 10^5^(dotted curves). In panel b, strains are coloured as in Fig. 1a. From numerical solutions of eqn. 1, with mutation *µ* = 10^−6^.

It is important to note that anomalous speeds require mutation to be non-zero, but persist even when the mutation rates between strains are vanishingly small (i.e. in the limit *µ →* 0, but still non-zero). This is surprising because, when the mutation rates are strictly zero, anomalous speeds do not occur and the strains invade independently at their monomorphic speeds (or not at all, if they are outcompeted in the stable equilibrium (Elliott and Cornell, 2012; Osnas et al., 2015)). However, a small amount of mutation, combined with exponential growth, is sufficient for the different strains to keep up with each other during the invasion and participate in the invasion. Other systems with anomalous invasion speeds are characterised by strongly co-operative interactions between strains, i.e. each strain has a strongly positive effect on the other (Weinberger et al., 2002, 2007). In our system, mutation is the only co-operative interaction between strains, so our case is unusual because anomalous invasion speeds exist even when cooperation (which is mutation, in this case) is vanishingly weak.

Furthermore, the limiting invasion speed when *µ →* 0 does not depend on the relative values *ν*_*i j*_ of the mutation rates, even if some mutation rates are zero (provided the system does not factorise into independent subsets of strains between which mutation is impossible). In particular, the invasion speed is the same for a system with “universal” mutation (*ν*_*ij*_ = 1 for all *i* and *j*) as for “nearest neighbour” mutation (*ν*_*ij*_ = 0 unless |*i - j*| = 1) (Fig. 1d, cyan curve in Fig. 1c). This means that our results should not only apply to species with a small set of discrete strains but also extend to the case of a very large set of strains where the traits can only mutate by small amounts at each generation.

This allows us to predict the invasion speed for a species with a continuous set of strains. We assume that each strain has a unique phenotype determined by *r* and *D*, and that there is a trade-off between *r* and *D* so that *r*(*D*) is a decreasing function of *D*. The invasion speed, after a sufficiently long time, will again be given by the largest permitted value of 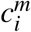 and 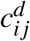 calculated using eqns. (6–8) for all strains, and pairs of strains of the species. It turns out that these equations have a geometric interpretation and that the invasion speed depends, in a simple way, on the shape of the *r*(*D*) curve. In particular, the existence of an anomalous invasion speed is determined by the curvature of the trade-off curve (Fig. 3, Supplementary Information Appendix S1).

**Figure 3:**
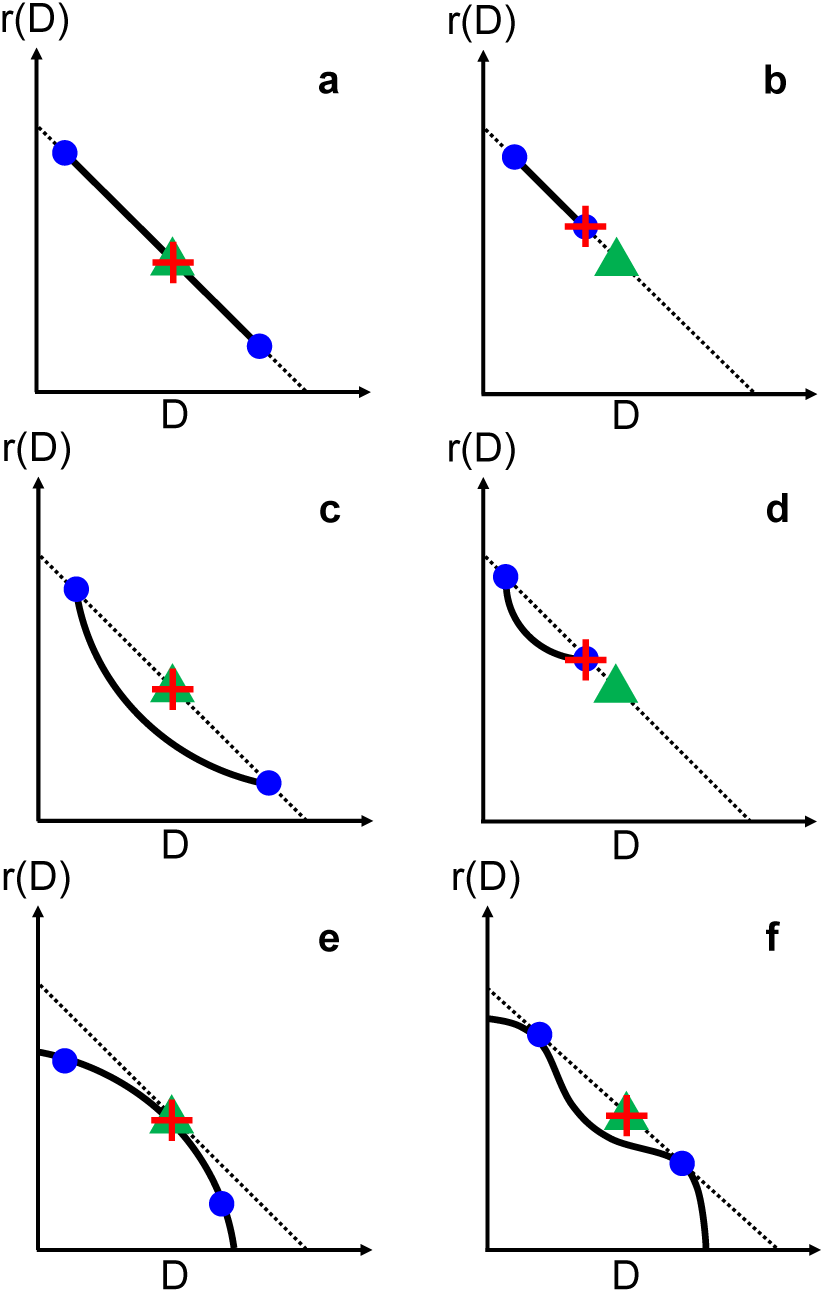
The geometry of the *D*-*r* trade-off determines which anomalous speeds, if any, give the eventual invasion speed for a species comprising a continuum of strains. Solid lines represent the trade-off curve between *D* and *r*. Dotted lines are chords that pass through points on the trade-off curve and terminate at the axes. Green triangles are the midpoint of these chords, and represents the “virtual strain” for any two strains that the chord passes through (see text). Blue points represent two particular strains of the species. The highest invasion speed in each case species is given by 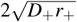, where *D*_+_ and *r*_+_ are evaluated at the red crosses. Panels a,b: for a straight-line trade-off, all pairs of species have the same chord and therefore the same virtual strain. In Panel a the virtual strain is a constituent of the species, but not in Panel b. In Panel c the virtual strain for the two extreme strains (blue circles) lies between them, and since the trade-off has positive curvature the virtual strain has a faster speed than any single constituent. In Panels b and d the virtual strain does not lie between any pair of points on the trade-off curve, so this does not yield a valid anomalous invasion speed. If the trade-off has negative curvature (Panel e) there are valid anomalous speeds, but none of their virtual strains lie above the trade-off so the asymptotic invasion speed is the fastest constituent monomorphic speed. In Panel F the chord is tangential to the trade-off at the two blue points, and since the whole trade-off curve lies below this chord the corresponding anomalous speed is faster than any other monomorphic or dimorphic speed for this species. This shows that the vanguard strains are not necessarily the ones with the highest values of *r* or *D*. We show further in the Methods section that the vanguard strains do not necessarily have the highest value of *rD* either.

Some simple algebra (Supplementary Information Appendix S1) shows that the dimorphic speed for two strains is equal to the monomorphic speed for a “virtual” strain, which lies at the midpoint (green triangles in Fig. 3) of the straight-line segment joining the two axes (dotted line in Fig. 3) and passing through the points representing the two strains (blue circles in Fig. 3) in *rD*-*r* space. Further simple algebra shows that, if the virtual strain lies between the two real strains, then conditions (8) are met and this is a valid dimorphic speed (Figs. 3a,c,e,f); otherwise, this is not a valid dimorphic speed for the species.

Therefore, if the trade-off curve is a straight line (Fig. 3a,b), then all pairs of strains have the same virtual strain (and therefore the same dimorphic speed) whether or not the trade-off curve encompasses this virtual strain, when the fastest possible speed will be the fastest monomorphic speed among the constituent strains. On the other hand, if the trade-off curve has negative curvature, Fig. (3e), then the virtual strain for any pair of strains either lies below the trade-off curve, or does not lie between the two real strains. In that case, none of the valid anomalous speeds are faster than the fastest monomorphic speed for the species. In both of these cases, the ultimate invasion speed will be the same as the monomorphic speed for the fastest strain in isolation, i.e. the strain with the highest value of *rD*(*r*) (see eqn. (6)).

However, if the trade-off curve has positive curvature (Fig. 3c,d), then the line segment joining the two most extreme strains will lie above the trade-off curve. If the range of *r* and *D* values is wide enough that the corresponding virtual strain lies between these extreme strains (Fig. 3c), then this will have the fastest dimorphic speed, which will be faster than any of the constituent monomorphic speeds. However if the range of *r* and *D* values are not wide enough (Fig. 3d), the species will invade at the fastest monomorphic speed. In the former case, the invasion will follow the anomalous speed when the vanguard strains have the highest *r* and the highest *D*. A further possibility is that the curvature of the trade-off curve is positive in some places and negative in others (Fig. 3f), for example if the trade-off is more acute at more extreme values. In this case, the vanguard strains will be the ones where the line joining them is tangential to the trade-off at both points, which will not represent the most extreme traits in the species (similarly to what was found in Fig. 1 for a species with a discrete set of trait values, where the vanguard strains can be found from Fig. 1a using the same construction as Fig 3f).

While the deterministic model, eqn. (1), predicts anomalous speeds for vanishingly small mutation, it has been shown for *N* = 2 that demographic stochasticity suppresses anomalous speeds when mutation or populations are small (Elliott and Cornell, 2013). We also find this to be the case for *N >* 2. While Fig. 4 b shows that a 5-strain species (black data) always invades at a speed close to that for a 2-strain species consisting of the vanguard strains (magenta data), this is only faster than the fastest monomorphic speed (blue data) if either the mutation rate is high or local populations (which are proportional to 1*/a*) are large. We again find that the vanguard strains (labelled 7 and 9 in Fig. 4a) need not be the most dispersive or the most fecund if the curvature of the dispersal-fitness tradeoff resembles Fig. 3f.

**Figure 4:**
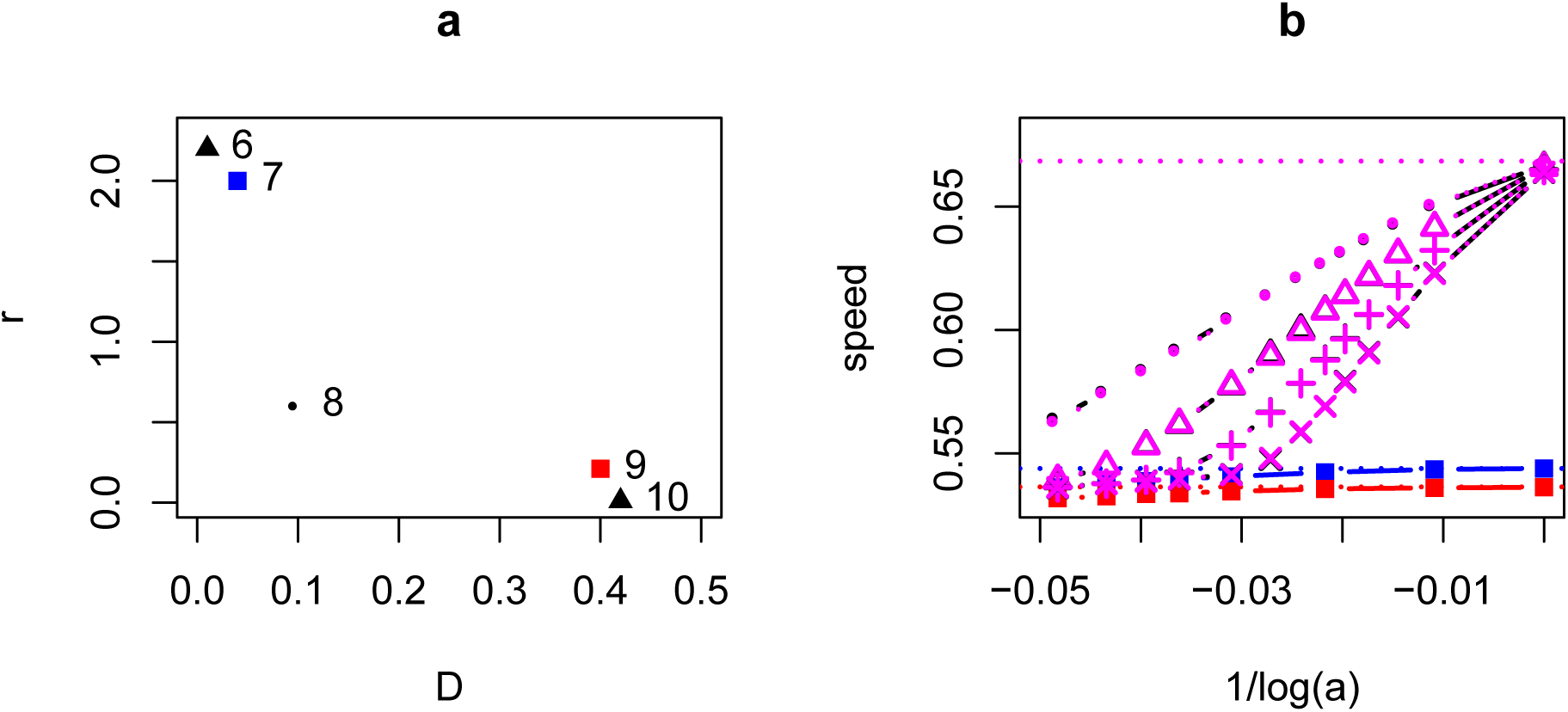
Effect of demographic stochasticity on anomalous invasion speeds. Panel a: trait values for 5 strains, where (*D*_*i*_, *r*_*i*_) = (0.01, 2.2), (0.04, 2),(0.1, 0.6), (0.4, 0.21), and (0.42, 0.01) for *i* = 6, 7, 8, 9, and 10 respectively. Strains 7 (the establisher, blue) and 9 (the disperser, red) are the vanguard strains. Panel b: invasion speeds as a function of the strength *a* of density dependence (which is inversely proportional to the carrying capacity) for species consisting of different combinations of the strains in Panel a. Red: strain 9 only; Blue: strain 7 only; magenta dotted: strains 7 and 9 only; black (partly obscured by the magenta data): all 5 strains together, with universal mutation. For the polymorphic species, different symbols denote the mutation rate: 10^−3^(circles), 10^−4^(triangles), 10^−5^(plus signs), 10^−6^ (multiplication signs). Horizontal dotted lines are predicted speeds (see Supplementary Information Appendix S1) in the deterministic limit *a* → 0 for the polymorphic species (magenta), strain 7 alone (blue), and strain 9 alone (red). From simulations of the stochastic discrete model.

## Discussion

We have presented a general theory for invasions by a species where dispersal ability varies among lineages and trades off against fitness. In some cases, invasions advance at the same rate as if the species consisted solely of the single strain that has the highest monomorphic speed. However, if the dispersal-growth curve has positive curvature, and extends over a wide enough range of dispersal rates, then an anomalous invasion occurs which is faster than that for any single constituent strain. This speed depends only on the traits of two “vanguard” strains, which need not be the most dispersive nor the most fecund. The invasion speed is insensitive to the mutation rates between strains (provided they are non-zero) or the details of inter-strain competition, and anomalous speeds can even be caused by strains that are outcompeted at equilibrium (Fig 2). Even if there were strains with arbitrarily large population growth or dispersal ability (which is unrealistic), the shape of the tradeoff could prevent the invasion accelerating without limit (see Fig. 3). This suggests that accelerating invasions are a sign that the species has not evolved to the biological limits of the trade-off at the time of sampling.

Our results are counterintuitive in a number of respects. First, while synergy between strains can cause polymorphic species to behave differently from monomorphic species (Bolnick et al., 2011), it is surprising that the effect is still strong (Figs 1c, 2b) even when mutation — the process generating this synergy — is very weak. Second, since the species exploits the dispersal ability of one strain and the fecundity of another, it is surprising that the vanguard strains do not necessarily have the highest dispersal or reproductive ability. Third, if strains interact synergistically, then one might expect than more than two strains would contribute. However, anomalous speeds with these surprising features could arise in a wide variety of models. When the invasion dynamics are determined by the linearised behaviour at low density (so-called “pulled waves”, van Saarloos 2003; Stokes 1976; Lewis and Kareiva 1993), a similar analysis to ours can be applied and the ultimate invasion speed for low mutation rate will be given either by the minimum of the *λ/k* curve for one strain, or the intersection between the curves of two strains (see eqn. (2) and Supplementary Information Appendix S1). Thus, no matter how many traits trade off against each other, or whatever density dependence acts at higher density, we predict that anomalous invasion speeds at low mutation rate depend only on the traits of two species.

Our results show that dispersal evolution during range expansion is still more complex than previously thought. Evolving invasion fronts are not necessarily led by the most dispersive strains (Travis and Dytham, 2002; Fronhofer and Altermatt, 2015; Phillips et al., 2010; Shine et al., 2011), nor even by the one that would invade most quickly in isolation (Osnas et al., 2015; Phillips and Perkins, 2019; Deforet et al., 2019). Phillips and Perkins (2019) have introduced the concept of “spatio-temporal fitness” to explain why evolution during invasions selects for a combination of dispersal and reproductive fitness. They do not consider mutation and therefore predict that the product of dispersal and reproductive fitness (or, equivalently, monomorphic invasion speed) is maximised. It would be interesting to extend their arguments to include mutation, to find a more intuitive argument for how vanguard strains are determined. Our results resemble those of Laroche et al. (2016), who showed that evolution in metacommunities can lead to mixed dispersal strategies in the presence of tradeoffs. However, the promotion of the densities of the vanguard strains is rather different from the evolution of a mixed strategy. First, while one vanguard strain leads the invasion, the density of the other one can trail behind non-vanguard strains (Figs. 1b,d and 2b) so the species is not dominated by these two strains. Second, the vanguard strains do not represent competing but rather cooperative strategies. We do not expect that anomalous invasion dynamics can lead to evolutionary branching between the vanguard strains since mutation between the vanguard strains is essential.

The anomalous dynamics we report require a combination of dispersal polymorphism, dispersal-fitness trade-off, and mutation during invasions. Without the trade-off, i.e. if all strains have the same population growth rate (Dockery et al., 1998; Bouin and Calvez, 2014), the invasion accelerates to the monomorphic speed of the most dispersive strain. Mutation is necessary because otherwise the strain with the fastest monomorphic speed leaves the others behind (Elliott and Cornell, 2012; Osnas et al., 2015). However, our results contrast with other studies on the effect of mutation on invasion dynamics, because anomalous invasion dynamics do not converge to monomorphic dynamics when the mutation rate becomes very small. Griette et al. (2015) computed the mutation-dependent invasion speed in a related 2-strain model, but without dispersal polymorphism so their invasion speeds converge to the fastest single-strain speed when the mutation rate approaches zero. It would also be possible to calculate, in our model, the first-order correction in the mutation rate *µ* to the invasion speed, but this will be much smaller than the effect of anomalous invasion speeds (a “zeroth order” effect) that we report here.

Our results predict that invasion dynamics depend critically on the curvature of the trade-off curve between dispersal and population growth rate, but while the existence of such trade-offs has been established (Hughes et al., 2003; Guerra, 2011; Karlsson and Johansson, 2008) their shape has not been quantified empirically in much detail. This trade-off arises from the organism diverting resources either to dispersal or to reproductive success, but the rates describing these abilities depend on the details of the organism’s anatomy and physiology so the trade-off curve could in principle take many different shapes. One plausible assumption would be that the organism’s reproductive success is proportional to the energy diverted to reproductive organs, and the distance of each dispersal event is proportional to the energy diverted to movement organs, which would imply a straight-line trade-off between fitness and dispersal distance. However, while population growth rate is directly proportional to fitness, the diffusion constant is proportional to the square of the dispersal distance, which would imply that the *D*(*r*) curve would be a parabola with positive curvature. On the other hand, diminishing returns would imply that an incremental improvement in one trait comes at a much greater cost when that trait is at the higher end of its range of possible values, which would suggest a trade-off with negative curvature (e.g. Fig. 3 e) or possibly a more complex curve such as in Fig. 3 f. Anomalous speeds could also occur when the trade-off curve has *negative* curvature, provided the species exists in distinct morphs (such as wing dimorphic crickets (Mole and Zera, 1993)) so that the trade-off is not a single continuous curve (and that the virtual strain with the fastest invasion speed lies between different segments of the trade-off curve). Thus, while anomalous speeds can be expected in a wide range of scenarios, to predict the species in which they occur we would need more detailed measurements of the trade-offs between dispersal and reproductive fitness than are currently available. Realistically, the trade-off curve represents the boundary of strains in r-D trait-space that would be produced from a collection of measurements from individuals.

Our models may be simple and generic, but we expect the predictions to apply in a wide range of scenarios. We used Brownian motion to model dispersal (i.e. assuming dispersal comprises many very short steps), but the analysis and our predictions would be similar if dispersal followed a jump process with thin-tailed (exponentially bounded) dispersal kernels. Our theory should apply to modestly fat-tailed) dispersal kernels of the class which lead to finite speed waves (i.e., those which are not exponentially bounded but have finite second moment), but, at first sight, not to “fatter” tailed kernels used to describe dispersal combining local movement and long-distance events (Clark et al., 1998; Clark, 1998; Kot et al., 1996).

Such kernels have been shown to predict accelerating invasions rather than a travelling wave of constant speed. However, it has been shown that fat-tailed dispersal kernels can describe a population of individuals, each performing Brownian motion but with a distribution of diffusion constants (Petrovskii et al., 2008). This is analogous to the dispersal polymorphism we discuss, but for the case of no tradeoff between *r* and *D*. In that case, and (unrealistically, but in common with Petrovskii et al. (2008)) assuming no upper limit to *D*, we also would predict invasions that accelerate without limit. However, if the more dispersive strains have a lower population growth rate, then our theory predicts that the invasion could follow an asymptotic constant invasion speed determined by the shape of this trade-off. This shows that it could be misleading to characterise a species by a single dispersal kernel, without considering whether intrinsic dispersal ability or reproductive ability might differ among individuals. We expect a more complex theory is needed to account for other factors such as non-linear diffusion or landscape heterogeneity, but our results constitute the first steps in this direction.

Our model assumes a haploid species, but we predict our theory to hold in any species where offspring inherit their strain (with rare mutation) from only one parent when population densities are low. The theory should apply for microbes as well as self-fertile plants, but cannot be simply extended to obligate sexually reproducing species with recombination where the population dynamics at low densities are non-linear. However, since anomalous speeds require mutation between strains in the low-density leading edge of the invasion, we find that they only arise in stochastic models if mutation or population sizes are large enough. For example, for the parameters in fig. 4b anomalous speeds require populations in excess of 10^10^ for a mutation rate of 10^−4^. We were unable to find a way to predict how large populations or mutation would need to be for a particular set of dispersal and growth parameters, as existing analytical methods for stochastic invasions (Brunet and Derrida, 1997; Hallatschek, 2011) cannot obviously be extended to our case, so we do not know whether there are cases where these numbers could be significantly lower. However, we expect that anomalous invasion dynamics could occur in viruses or microbial systems, particularly infectious diseases, where hosts constitute demes within a large local population. Indeed, anomalous invasion speeds have been shown to occur in SIR-type systems that allow “virus shedding” which allow the production of virions proportional to infected agents (Reluga, 2004). RNA viruses can have mutation rates as high as 10^−3^-10^−5^ per base pair per generation (Drake et al., 1998), and can exist at densities of 10^12^ individuals per gramme of human faeces (Atmar et al., 2008), so within-host populations could easily be high enough for anomalous invasion speeds. It is possible that anomalous speeds could occur in species with smaller populations when diversity is maintained at the invasion front by processes other than mutation. This might arise due to Allee effects (Roques et al., 2012) or sexual reproduction, for example, but in those cases our analytical methods could not be applied because travelling waves would be “pushed” rather than “pulled” (van Saarloos, 2003).

Another much-studied aspect of evolution during invasions is “expansion load” (Peischl et al., 2013), a progressive decrease in fitness and/or the speed of invasion or range expansion (Peischl et al., 2015) due to the fixation of deleterious mutations near the invasion front. Evolution of dispersal can act in combination and remove the deceleration caused by expansion load (Peischl and Gilbert, 2018). However, it is not clear that anomalous invasion speeds can occur in populations where expansion load is significant. This is because the former requires large population sizes in the leading edge of the front, but the latter required populations small enough for deleterious mutations to become fixed. These two phenomena therefore represent complementary potential outcomes for evolution during range expansions.

Many global threats are due to biological invasions: food security from crop pests; antibiotic resistance; local disease outbreaks; and loss of biodiversity. Many other important phenomena can be modelled with similar equations, e.g. the cell population dynamics of tumor growth. Our results show that it is not safe to model an invading polymorphic population with a single “average” strain, or as multiple strains that vary in a single trait. Nevertheless, we show that the invasion speed can be straightforwardly predicted from the dispersal-fitness trade-off alone. Our results highlight the importance of understanding and quantifying such trade-offs, as well as the need to account for evolution when considering central questions in ecology.

## Acknowledgments

VAK was supported by a PhD studentship from the Natural Environment Research Council (NERC ACCE DTP, grant number NE/L002450/1). We would like to thank Mike Begon, Andy Fenton, Ilik Saccheri, and Ben Phillips for helpful comments on this manuscript. We would like to thank the Plant Health Centre Scotland for allowing VAK time to complete edits of this manuscript outside of his usual role.

