## Supporting Information 1 for "Anomalous invasion dynamics due to dispersal polymorphism and dispersal-reproduction trade-offs"

### Appendix S1: supplementary methods

#### Mathematical analysis

We consider an asexual haploid species consisting of $N$ strains (i.e. genotypes), where each strain has a distinct growth rate and dispersal phenotype, with the possibility of mutation between strains at birth (derivation can be found in Elliott and Cornell (2012) supplementary material). We assume density-dependent competition between strains and that dispersal can be approximated by diffusion, and model the dynamics using the following spatial Lotka-Volterra model

$$\frac{\partial n_i}{\partial t} = \overbrace{D_i \frac{\partial^2 n_i}{\partial x^2}}^{\text{Dispersal}} + \overbrace{r_i n_i (1 - \sum_{j \in S} C_{ji} n_j)}^{\text{Lotka-Voterra dynamics}} + \tag{S.1}$$

$$\overbrace{+ \sum_{j \in S \setminus \{i\}} \mu_{ji} n_j - \sum_{j \in S \setminus \{i\}} \mu_{ij} n_i}^{\substack{\text{Mutation dynamics from population i} \\ \text{into populations } j \text{ and vice-versa}}}, \quad \forall i \in S,$$

where $S = \{1, 2, ..., N\}$ is the set of all strains, $D_i$ and $r_i$ are respectively the diffusion constant and population growth rate of the $i$th strain, $n_i$ is the density of the $i$th strain, $C_{ji}$ is the competitive effect of strain $j$ on $i$, and $\mu_{ij}$ the mutation rate from strain $i$ to strain $j$. This equation is derived in a similar way to Elliott and Cornell (2012), by assuming that the per capita birth rate for strain $i$ is $b_i$, that a small fraction $\frac{\mu_{ij}}{b_i}$ of these births mutate into strain $j$, and that the death rate is $d_i + r_i \sum_{j \in S} C_{ji} n_j$, where $r_i = b_i - (\sum_{j \in S} \mu_{ij}) - d_i$ . All parameters are assumed to be positive in order to be biologically relevant or, in the case of $C_{ji}$, so that intraspecific interactions are competitive rather than cooperative.

We considered a population in which all strains initially coexist in the semi-infinite landscape $x < 0$ and subsequently invade unoccupied habitat $x > 0$. If any strain $i$ were present in the landscape in isolation (which would require $\mu_{ij} = 0$) the classic result for the predicted invasion speed $c_{mi} = 2\sqrt{D_i r_i}$ holds (Elliott and Cornell, 2012; Fisher, 1937). From here on, we discuss only the case when all strains coexist (which requires that at least some of the mutation rates are non-zero).

The general system of spatially uniform equations has a trivial unstable equilibrium point $\mathbf{n}^*_{trivial} = \mathbf{0}$ and a non-trivial stable equilibrium $\mathbf{n}^*_{non-trivial} \in \mathbb{R}^N$. It has recently been proven (Girardin, 2018) that the solutions to equations of the form eqn. (S.1) under these initial conditions are travelling waves of the form $n_i = N_i(x - c^* t)$, where $c^*$ is obtained by the solution of an eigenvalue problem (previously, except in a few special cases, it had only been proven that the value of $c^*$ obtained in this way was a lower bound for the spreading speed).

Linearising eqn. (S.1) around the unstable equilibrium point $\mathbf{n}^*_{trivial} = \mathbf{0}$ leads to the form

$$\frac{\partial n_i}{\partial t} = D_i \frac{\partial^2 n_i}{\partial x^2} + \tilde{r}_i n_i + \sum_{j \in S \setminus \{i\}} \mu \nu_{ji} n_j, \quad \forall i \in S, \tag{S.2}$$

where $\tilde{r}_i = r_i - \sum_{j \in S \setminus \{i\}}^N \mu_{ij}$ and we have defined $\nu_{ij} = \frac{\mu_{ij}}{\mu}$ so that we can take the limit $\mu \to 0$ while keeping $\nu_{ij} = O(1)$. Using Theorem 1.7 of Girardin (2018), the spreading speed for eqn. (S.1) is given by

$$c^* = \min_k \max_i \left( \frac{\lambda_i(k, \mu)}{k} \right) \tag{S.3}$$

where $\lambda_i(k, \mu)$, $i \in S$ are the eigenvalues of the matrix

$$\mathrm{M}(k, \mu) = \begin{bmatrix} D_1 k^2 + \tilde{r}_1 & \mu\nu_{21} & \cdots & \mu\nu_{N1} \\ \mu\nu_{12} & D_2 k^2 + \tilde{r}_2 & \cdots & \mu\nu_{N2} \\ \vdots & \vdots & \ddots & \vdots \\ \mu\nu_{1N} & \mu\nu_{2N} & \cdots & D_N k^2 + \tilde{r}_N \end{bmatrix}.$$

While the eigenvalues of this $N \times N$ matrix cannot in general be computed in closed form, we can proceed by making the biologically realistic assumption that mutation is a rare event, so that $\mu \ll 1$. The invasion speed will then be well approximated by the limit of the invasion speed as $\mu \to 0$. When $\mu = 0$ all of the off diagonal elements are zero so the eigenvalues equal

$$\lambda_i(k, 0) = D_i k^2 + r_i, \quad i \in S$$

(since $\lim_{\mu \to 0} \tilde{r}_i = r_i$).

Taking the limit $\mu \to 0$ in eq. (S.3)

$$\begin{aligned} \lim_{\mu \to 0} c^* &= \lim_{\mu \to 0} \min_k \max_i \left( \frac{\lambda_i(k, \mu)}{k} \right) \\ &= \min_k \max_i \left( \frac{\lambda_i(k, 0)}{k} \right), \end{aligned}$$

In the second line of the preceeding equation, we have swapped the order of the limit $\mu \to 0$ with the minimum and maximum; this is justified because the eigenvalues of a matrix depend continuously on its elements, so the eigenvalues of $M$ converge uniformly to $\lambda_i(k,\mu)$ as $\mu \to 0$. We can therefore find the limiting value of the invasion speed $c^*$ as $\mu \to 0$ by plotting the curves $D_ik + \frac{r_i}{k}$ as a function of $k$, selecting the upper envelope of these curves, and then finding the value of $k$ for the lowest point on this upper envelope. This procedure is illustrated in Figure S.1.

The invasion speed can be given by the minima of one of the $D_ik + \frac{r_i}{k}$ curves (e.g. as in top row, fig. S.1), i.e.

$$c_i^m = 2\sqrt{D_i r_i},$$

which is the speed at which a species comprising strain $i$ in isolation would invade. Alternatively, it can be given where the curves $D_ik + \frac{r_i}{k}$ for two strains (say, $l$ and $m$) cross, which occurs when $k = k^*$ where

$$D_l k^* + \frac{r_l}{k^*} = D_m k^* + \frac{r_m}{k^*} \tag{S.4}$$

$$\Rightarrow k^{*2} = \frac{r_m - r_l}{D_l - D_m}, \tag{S.5}$$

and therefore at a speed

$$c_{lm}^d = D_l k^* + \frac{r_m}{k^*} = \pm \frac{r_l D_m - r_m D_l}{\sqrt{(r_l - r_m)(D_m - D_l)}} \tag{S.6}$$

Note that, for $k$ in eq. (S.5) to be real, we require that the strain that has the higher value of $D$ also has the lower value of $r$, i.e. there is a tradeoff between dispersal and reproductive fitness. This point can only be a minimum of $\max_i \left(\frac{\lambda_i(k,0)}{k}\right)$ if the two $D_lk + \frac{r_l}{k}$ curves have gradients with opposite signs (see Fig. S.1 bottom right), i.e. if

$$\left(D_l - \frac{r_l}{k^{*2}}\right)\left(D_m - \frac{r_m}{k^{*2}}\right) < 0$$

$$\Rightarrow \frac{r_l r_m D_l D_m}{(r_l - r_m)^2}\left(2 - \frac{r_l}{r_m} + \frac{D_l}{D_m}\right)\left(2 - \frac{r_m}{r_l} + \frac{D_m}{D_l}\right) > 0,$$

which requires either that

$$\frac{r_l}{r_m} + \frac{D_l}{D_m} < 2 \quad \text{and} \quad \frac{r_m}{r_l} + \frac{D_m}{D_l} < 2 \tag{S.7}$$

or

$$\frac{r_l}{r_m} + \frac{D_l}{D_m} > 2 \quad \text{and} \quad \frac{r_m}{r_l} + \frac{D_m}{D_l} > 2. \tag{S.8}$$

However, it is straightforward to show that eqn. (S.7) cannot be satisfied when $r_i \geqslant 0$, and $D_i \geqslant 0$, $\forall i \in S$: that would require both $\frac{D_l}{D_m} < 2 - \frac{r_l}{r_m}$ , implying

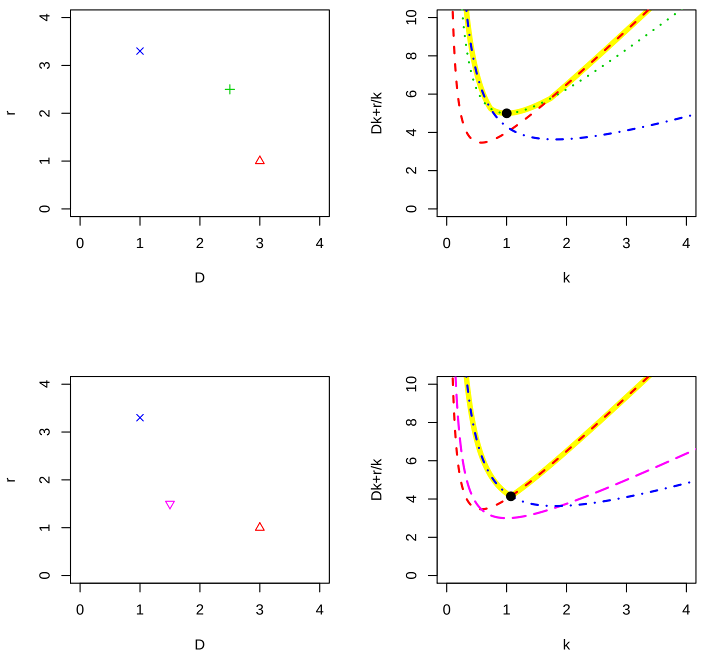


Figure S.1: Finding the invasion speed for two different 3-strain species where the speed is not anomalous (top row) and is anomalous (bottom row). Panels on the left illustrate the phenotypic traits of the constituent strains, and on the right are graphs of the numerically computed value of $\frac{\lambda_P(k,\mu)}{k}$ (solid yellow line) and the functions $\frac{\lambda_i(k,0)}{k}$ (other curves). The mutation rate is small ($\mu = 0.001$) so the $\frac{\lambda_P(k,\mu)}{k}$ is close to the upper branch of the $\frac{\lambda_i(k,0)}{k}$ curves. The wave speed is given by the lowest value of the yellow curve (black solid circle). When $\mu \to 0$, the wave speed is either given by the minima of one of the $\frac{\lambda_i(k,0)}{k}$ (top row), or it is given by the point where two of the $\frac{\lambda_i(k,0)}{k}$ curves cross (bottom row). In the first case (top row), the invasion speed is not anomalous, because it is given by the minima of the $\lambda_i/k$ curves for one of the strains, and is therefore the same as the monomorphic speed for that strain. In the second case (bottom row), the invasion speed is anomalous because the black circle lies above the minima of all of the single-strain $\lambda_i/k$ curves. In the rop row the species comprises 1, 2, and 3, and in the bottom row strains 1, 3, and 4, where the strains' phenotypic traits are: $(D_1, r_1) = (3, 1)$ (red upward triangle or short dashed curve); $(D_2, r_2) = (2.5, 2.5)$ (green plus sign or dotted line); $(D_3, r_3) = (3, 1)$ (blue multiplication sign or dash-dottted line); $(D_4, r_4) = (1.5, 1.5)$ (magenta downward triangle or long-dashed curve).

$\frac{r_l}{r_m} < 2$, and also $\frac{D_m}{D_l} < 2 - \frac{r_m}{r_l}$, implying $\frac{r_l}{r_m} > 2$. Therefore, (S.8) must be met for $c^d_{lm}$ to be a valid wave speed for the system. Note that the same condition was derived for the two phenotype case by Elliott and Cornell (2012).

Therefore, the invasion speed follows eqns. (6–8) in the main paper.

### Numerical simulations

We solve eqns. (1) numerically in R v. 3.5.3 (R Core Team, 2013) using the ode.1D() function from the deSolve package (v. 1.24) (Soetaert et al., 2010), placing the model on a 1 dimensional lattice and replacing the Laplacian with a second order spatial difference. Reflecting boundaries were used. These solutions all produced travelling wave solutions that rapidly converged to constant speeds. The position of the front was defined by using exponential interpolation to estimate the value of $x$ where the population density would be 0.001, and the wave speed estimated by differences in front position over 100 time steps. Lattices of 30000 or 40000 sites were used (large enough so that the front never reached the end of the system). To avoid spurious results due to roundoff errors when densities were close to the floating point precision of the computer, the densities were at all times set to zero if they were less than $10^{-300}$. R code is available at Figshare DOI:XXX.(Soetaert et al., 2010)

### Geometric interpretation of anomalous speeds

A straight line passing through points representing strains $i$ and $j$ in $(D, r)$ space has equation

$$r = r_i + (D - D_i)\frac{r_j - r_i}{D_j - D_i}, \tag{S.9}$$

so that it meets the axes at the points $(D, r) = \left(\frac{r_i D_j - r_j D_i}{r_j - r_i}, 0\right)$ and $(D, r) = \left(0, \frac{r_i D_j - r_j D_i}{D_j - D_i}\right)$. The midpoint of the line segment joining the two axes is then

$$(D, r) = (D_{vij}, r_{vij}) = \left(\frac{r_i D_j - r_j D_i}{2(r_j - r_i)}, \frac{r_i D_j - r_j D_i}{2(D_j - D_i)}\right), \tag{S.10}$$

and the monomorphic speed for this "virtual" strain is then $c = 2\sqrt{r_{vij} D_{vij}}$, which gives the same expression as $c^d_{ij}$ from eq. (7) of the main paper. Thus, the dimorphic invasion speed for strains $i$ and $j$ is the same as the monomorphic speed for this virtual strain.

We assume (without loss of generality) that $r_i < r_j$, which implies that $D_j < D_i$ in order for the dimorphic speed in eqn. (7) of the main paper to be real. The condition for the virtual strain to lie between the real strains is then

$$r_i \leq r_{vij} \leq r_j,$$

and it is straightforward to show that this leads to exactly the same conditions as eqns. (8) in the main paper. it is also straightforward to show that the virtual

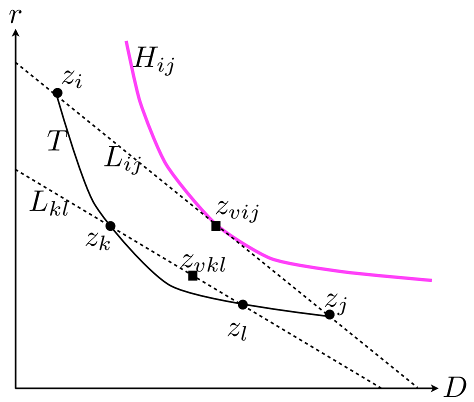


Figure S.2: The invasion speed when $D-r$ trade-off curve (represented by $T$) has positive curvature is the dimorphic speed for the virtual strain $v_{ij}$ (represented by a point $z_{vij}$) for the two most extreme strains $i$ and $j$ (represented by points $z_i$ and $z_j$), provided $z_{vij}$ lies between $z_i$ and $z_j$. $H_{ij}$ is a hyperbola $r = \frac{D_{vij}r_{vij}}{D}$ that represents the locus of strains that would have the same monomorphic speed as $v_{ij}$; all strains represented by points below $H_{ij}$ have $Dr < D_{vij}r_{vij}$, and therefore slower monomorphic speeds than the virtual strain $v_{ij}$. $L_{ij}$ (resp $L_{kl}$) is a straight line segment passing through $z_i$ and $z_j$ (resp. $z_k$ and $z_l$) and terminating on the $r$ and $D$ axes. As shown above, $z_{vij}$ (resp. $z_{vkl}$) lies at the midpoint of $L_{ij}$ (resp. $L_{kl}$). Since $H_{ij}$ is tangential to $L_{ij}$ at $z_{vij}$, the monomorphic speed for the virtual strain $z_{vij}$ is faster than the monomorphic speed for any other strain on $T$. Moreover, since the virtual strain for any pair of strains $k$ and $l$ must lie between $z_k$ and $z_l$ for the corresponding dimorphic speed to be a valid invasion speed, so $z_{vkl}$ must lie below $L_{ij}$ and therefore below $T$, no dimorphic speed $c^d_{kl}$ can exceed $c^d_{ij}$.

strain has the highest value of $rD$, and therefore the highest monomorphic speed $2\sqrt{rD}$, of any strain on the line segment defined by eqn. (S.9). This implies that the hyperbola

$$r = \frac{D_{vij}r_{vij}}{D}.$$

along which the monomorphic speed equals $c^d_{ij}$, is tangential to the line segment at $(D, r) = (D_{vij}, r_{vij})$

These geometric constructions allow us to find the invasion speed from the geometric properties of the trade-off curve alone. As shown in fig. S.2, if the curve segment representing the trade-off has positive curvature then the invasion speed is the anomalous dimorphic speed for the two most extreme strains, provided the corresponding virtual strain is bracketed by these extreme strains (i.e. that the condition eqn. (4) is satisfied so that this is a valid invasion speed). A similar construction shows that, if the trade-off curve has negative curvature, then no valid dimorphic speed is faster than the fastest monomorphic speed on the trade-off curve.

If the trade-off curve has segments with both positive and negative curvature (e.g. Figure 4 f) then the pair of strains that give the fastest dimorphic speed need not be at the extrema of the trade-off curve $r = r(D)$ provided there is a pair of strains with diffusion constants $D_1$ and $D_2$ where $\frac{dc_{12}^d}{dD_1} = 0 = \frac{dc_{12}^d}{dD_2}$. Taking derivatives of eqn. (3) gives

$$\frac{dc_{12}^d}{dD_1} = \frac{\{D_1 r(D_2) + D_2 r(D_1) - 2D_2 r(D_2)\}\left[(D_1 - D_2) r'(D_1) + r_2 - r_1\right]}{2\left((D(r_1) - D(r_2))(D_2 - D_1)\right)^{3/2}} = 0,$$
$$\frac{dc_{12}^d}{dD_2} = \frac{\{D_1 r(D_2) + D_2 r(D_1) - 2D_1 r(D_1)\}\left[(D_1 - D_2) r'(D_2) + r_2 - r_1\right]}{2\left((D(r_1) - D(r_2))(D_2 - D_1)\right)^{3/2}} = 0.$$

If either of the terms in braces is zero, then as seen in eqn. (S.10), the virtual strain is identical to either strain 1 or strain 2, so the dimorphic speed is the same as the monomorphic strains. If the dimorphic speed is truly anomalous, i.e. strictly greater than any monomorphic speed, then the terms in braces must be non-zero, so we must have

$$r'(D_1) = r'(D_2) = \frac{r(D_1) - r(D_2)}{D_1 - D_2}.$$

The term on the right hand side is equal to the gradient of the straight line joining the two strains, so this line must therefore be tangential to the trade-off curve at both points (see Fig. 4 f).

To show that the vanguard morphs need not have the highest values of $rD$ on the trade-off curve, we provide an illustrative example in figure S.3

### Discrete model

To test the robustness of our results to stochasticity, we define a model discrete in space and time, based on multi-strain Beverton-Holt dynamics. Let $n_i(x,t)$ be the density of strain $i$ on site $x$ (on a 1-dimensional lattice) at integer time step $t$. We assume that, at each time step, density-independent birth is followed by density-dependent death and then dispersal to neighbouring sites. After birth, assuming fecundity $r_i$ and mutation probability $\mu$ per birth, the expected number of individuals of strain $i$ is

$$m_i(x,t) = n_i(x,t)\left(1 + r_i(1 - N\mu)\right) + \mu \sum_{j \in S} r_j n_j(x,t),$$

and density-dependence causes a fraction $p_i = \frac{1}{1 + a\sum_j m_i}$ to die. We implement stochasticity by combining these steps and using a Poisson pseudorandom number generator, so that after births and deaths the number of survivors is

$$\tilde{m}_i(x,t) \sim \mathrm{Pois}(m_i(x,t)).$$

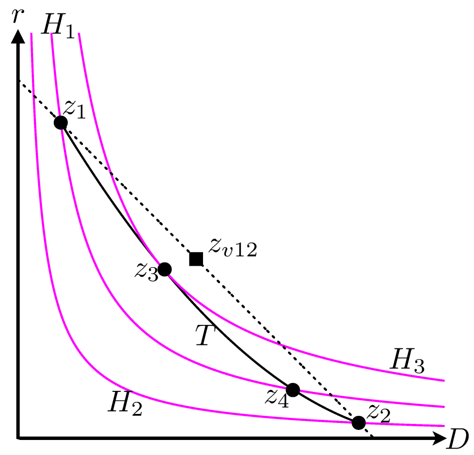


Figure S.3: Example to illustrate that vanguard strains do not necessarily have the highest value of $rD$ on the trade-off curve. A convex tradeoff curve (solid black line $T$) in $(D, r)$ space has been constructed such that there is a hyperbola $H_3$ that is tangential to it at a point $z_3$, which is not one of $T$'s endpoints ($z_1$ and $z_2$). In addition, the curve has been constructed such that the point $z_{v12}$ representing the virtual strain (constructed as in fig. S.2) lies between $z_1$ and $z_2$, so this tradeoff curve gives rise to an anomalous invasion speed. Any curve $rD = C_1$ lies wholly above a curve $rD = C_2$ provided $C_1 > C_2$. In particular, the hyperbole $H_1$ and $H_2$, passing respectively through $z_1$ and $z_2$, lie below $H_3$, so the value of $rD$ is higher at $z_3$ than at either $z_1$ or $z_2$ (or, for that matter, any other point on $T$).

In the dispersal step, a fraction $D_i$ disperses to either neighbouring site. We implement this in two stages: first generate the number of dispersers using a Binomial Pseudorandom number generator:

$$\tilde{d}_i(x,t) \sim \text{Bin}(\tilde{m}_i(x,t), 2D_i),$$

then split these into individuals moving right ($\tilde{d}_i^+$) or left ($\tilde{d}_i^-$):

$$\tilde{d}_i^+(x,t) \sim \text{Bin}(\tilde{d}_i(x,t), D_i)$$
$$\tilde{d}_i^-(x,t) = \tilde{d}_i(x,t) - \tilde{d}_i^+(x,t)$$

so that the densities at the beginning of the next timestep are

$$n_i(x,t+1) = \tilde{m}_i(x,t) - \tilde{d}_i(x,t) + \tilde{d}_i^{\,+}(x-1,t) + \tilde{d}_i^{\,-}(x+1,t).$$

We implemented this model in R v. 3.5.3 (code available upon request. In the limit when $a$ is very large, stochastic fluctuations become unimportant and the model is equivalent to the deterministic model given by eqns. (3-5) in the main paper. We can predict the wave speed in this deterministic limit by linearising around the low density limit, which yields the linearised equations:

$$n_i(x,t+1) = \bar{n}_i(x,t)(1-2D_i) + D_i(\bar{n}_i(x-1,t) + \bar{n}_i(x+1,t)),$$

where

$$\bar{n}_i(x,t) = n_i(x,t)(1 + r_i(1-N\mu)) + \mu \sum_{j \in S} r_j n_j(x,t).$$

Making the ansatz $n_i \sim \lambda^t \omega^x$,we find (in a similar way to the PDE model) that the eigenvalues $\omega$ satisfy

$$\omega_i(\lambda,\mu) = (1+r_i)\left\{1 + D_i(\lambda + \lambda^{-1} - 2\right\} + O(\mu).$$

We can therefore calculate possible wave speeds as a function of $\lambda$ as

$$c_i(\lambda,\mu) = \frac{\log \omega_i(\lambda,\mu)}{\log \lambda}. \tag{S.11}$$

By analogy with the PDE model, we hypothesise that the selected wavespeed is found by minimising over $\lambda$ the highest wave speed at each lambda. In the limit $\mu \to 0$, this can be achieved by

$$\lim_{\mu \to 0} c(\mu) = \min_{\lambda} \max_{i} c_i(\lambda,\mu).$$

This is illustrated for the strains used in Fig. 4 of the main paper in Supplementary Figure 4.

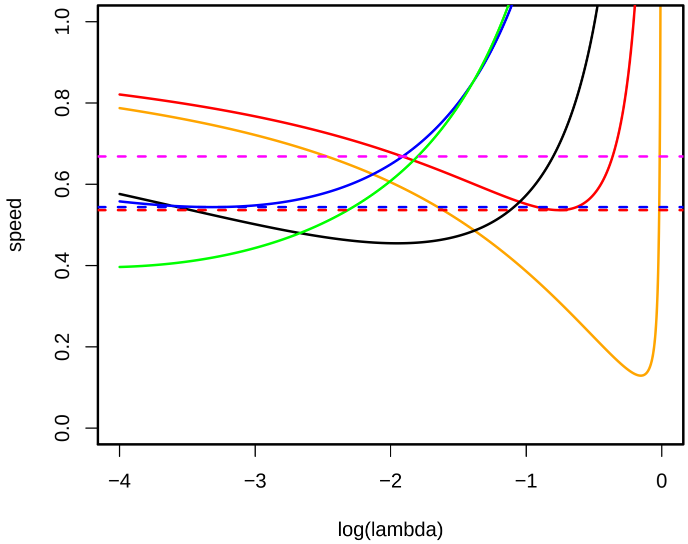


Figure S.4: Prediction of invasion speeds for discrete model, from eqn. (S.11). Strains are coloured as in Fig. 4a in the main paper. The minimum of the upper branch is given by the crossing of the blue and red curves, which corresponds to an anomalous speed 0.6685 generated by strains 7 and 9 (horizontal magenta dotted line). Horizontal blue and red curves give the predicted monomorphic speeds for strains 7 (0.5439) and strain 9 (0.5364) respectively.
